# Decoupling the molecular versatility of aminoglycosides via drug or target modification enables community-wide antiphage defense

**DOI:** 10.1101/2024.02.27.582341

**Authors:** Larissa Kever, Qian Zhang, Aël Hardy, Philipp Westhoff, Yi Yu, Julia Frunzke

## Abstract

The ongoing arms race between bacteria and phages has forced bacteria to evolve a sophisticated set of antiphage defense mechanisms that constitute the bacterial immune system. In our previous study, we highlighted the antiphage properties of aminoglycoside antibiotics, which are naturally secreted by *Streptomyces*. Successful inhibition of phage infection was achieved by addition of pure compounds and supernatants from a natural producer strain highlighting the potential for community-wide antiphage defense. However, given the dual functionality of these compounds, neighboring bacterial cells require resistance to the antibacterial activity of aminoglycosides to benefit from the protection they confer against phages. In this study, we demonstrated the successful uncoupling of antiphage and antibacterial properties via different aminoglycoside-resistance mechanisms encompassing drug and target site modifications. Furthermore, we confirmed the antiphage impact of aminoglycosides in a community context by co-culturing phage-susceptible, apramycin-resistant *S. venezuelae* with the apramycin-producing strain *Streptoalloteichus tenebrarius*. Given the prevalence of aminoglycoside resistance among natural bacterial isolates that allow functional uncoupling of these compounds, this study highlights the ecological relevance of chemical defense via aminoglycosides at the community level.

## Introduction

Owing to the constant threat of viral predation exerted by bacteriophages (or phages), bacteria have evolved an extensive repertoire of antiphage defense mechanisms. In recent years, advancements in high-throughput bioinformatics and experimental approaches have expanded our understanding of bacterial antiphage defense to over a hundred different systems, unveiling an unexpectedly rich portfolio of systems and mechanisms (Georjon & Bernheim, 2023). While previous knowledge of antiphage mechanisms was largely confined to restriction-modification and CRISPR-Cas systems targeting invading nucleic acids, recent discoveries have exposed remarkably intricate bacterial immune strategies. These include systems that rely on intracellular signal transduction facilitated by cyclic (oligo-)nucleotides and systems recognizing conserved structural patterns in viral proteins to initiate immune responses (Cohen *et al*., 2019, Ofir *et al*., 2021, Tal *et al*., 2021, Tal & Sorek, 2022, Gao *et al*., 2022). Recent studies also revealed several mechanisms that are shared among cells within a microbial community. These communal defences encompass the extrusion of membrane vesicles serving as a phage decoy, quorum sensing-based activation of defense systems, biofilm formation and secretion of antiphage small molecules (Luthe *et al*., 2023).

Environmental bacteria, particularly *Streptomyces* spp., are sophisticated producers of bioactive compounds endowed with antibacterial, antifungal or even anticancer properties (Barka *et al*., 2016). The natural producers usually possess self-resistance mechanisms to prevent auto-toxicity (Hopwood, 2007), thereby conferring them a fitness advantage over other microorganisms (Tyc *et al*., 2017). Shortly after their discovery, bacterial secondary metabolites emerged as a pivotal breakthrough to combat bacterial and fungal infections or applications in chemotherapy (Al-shaibani *et al*., 2021). Interestingly, recent studies indicate that bacterial small molecules also exhibit antiphage activity (Hardy *et al*., 2023). The *Streptomyces*-derived, naturally secreted anthracyclines doxo- and daunorubicin, which are primarily recognized as anticancer drugs, have been shown to efficiently inhibit infection of dsDNA phages, presumably by specifically intercalating into phage DNA (Kronheim *et al*., 2018). In our previous report, we expanded the repertoire of known antiphage molecule classes by demonstrating the antiphage properties of aminoglycoside antibiotics in widely divergent bacterial hosts (Kever *et al*., 2022a, Hardy *et al*., 2023).

Aminoglycosides are bactericidal antibiotics that inhibit protein translation by binding to the 16S rRNA of the 30S ribosomal subunit (Krause *et al*., 2016a). While different aminoglycosides exhibit varying specificity for distinct regions on the ribosomal A-site, they all induce a conformational change leading to the promotion of mistranslation by inducing codon misreading upon the delivery of aminoacyl transfer RNA. Despite the initial recognition of their antiphage properties as early as the 1960s (Schindler, 1964, Bowman, 1967, Brock *et al*., 1963), the exploration of these features in the context of bacterial antiphage immunity was not further pursued. In-depth analysis of this observation using aminoglycoside-resistant strains revealed a strong inhibition of phage infection by addition of pure apramycin as well culture supernatant (=spent medium) of the natural apramycin producer *Streptoalloteichus tenebrarius* (Tamura *et al*., 2008), hinting towards an ecological relevance of such chemically-mediated antiphage defense via aminoglycosides. Their secretion into the environment could create an antiviral microenvironment, thereby providing a chemical defense against phages at the community level (Hardy *et al*., 2023, Luthe *et al*., 2023). However, in order to benefit from this, neighboring cells or mycelial structures must possess resistance mechanisms to counteract the antibacterial impact of the compound without abolishing the antiphage effect. For aminoglycosides, three main resistance mechanisms are currently known including aminoglycoside-modifying enzymes (AME) for drug modification, 16S rRNA methyltransferases for target site modification and increased export or decreased uptake (Garneau-Tsodikova & Labby, 2016). Recent investigations reveal the widespread distribution of AMEs across different continents and terrestrial biomes, wherein ∼25% of sequenced bacteria carrying at least one AME (Pradier & Bedhomme, 2023). This distribution is expected to be closely correlated with the high mobility of AMEs. Remarkably, ∼40% of the detected AMEs are located on mobile genetic elements, facilitating their acquisition and broadening of the resistance spectrum via horizontal gene transfer (Pradier & Bedhomme, 2023). Additionally, they are often carried by phage plasmids, allowing them to be easily transferred across bacteria upon infection without direct cell-cell contact (Pfeifer *et al*., 2022).

In this study, we systematically examined the influence of various resistance mechanisms, encompassing a set of AMEs and a 16S rRNA methyltransferase, on the antiphage properties of aminoglycosides. Remarkably, our results demonstrated the effective uncoupling of antiphage and antibacterial properties through diverse drug modifications as well as target site modification across different aminoglycosides. This evidence shared here provides the pre-requisite for making a community-wide antiphage defense, even on an interspecies level, conceivable.

## Results

### Uncoupling of antiphage from antibacterial properties via drug and target site modification

Natural producers of secondary metabolites typically exhibit resistance to the antimicrobial molecules they synthesize (Hopwood, 2007, Tenconi & Rigali, 2018). This characteristic gains significance, particularly in the screening of small molecules for antiviral properties, as potential toxic effects on bacterial growth could mask any observed inhibition of phage infection. In a previous study, we showed the inhibition of phage infection for a broad range of aminoglycoside antibiotics by using strains resistant to the respective molecules (Kever *et al*., 2022a). In case of the used model aminoglycoside apramycin belonging to the subclass of aminoglycosides with mono-substituted 2-desoxystreptoamin (2-DOS) ring (Krause *et al*., 2016a), this resistance mechanism relied on the acetylation of the 3’ amino group of the 2-desoxystreptamin ring via aminoglycoside N(3)‐ acetyltransferase AAC(3)-IVa (Magalhaes & Blanchard, 2005) (Figure 1**A**). In this study, we aimed to investigate the impact of diverse bacterial (self-)resistance mechanisms on the antiphage properties of different aminoglycoside antibiotics. Our goal was to discern whether the observed decoupling of antibacterial and antiphage properties through resistance mechanisms is a pervasive trait among different mechanisms found in bacterial genomes.

**Figure 1:**
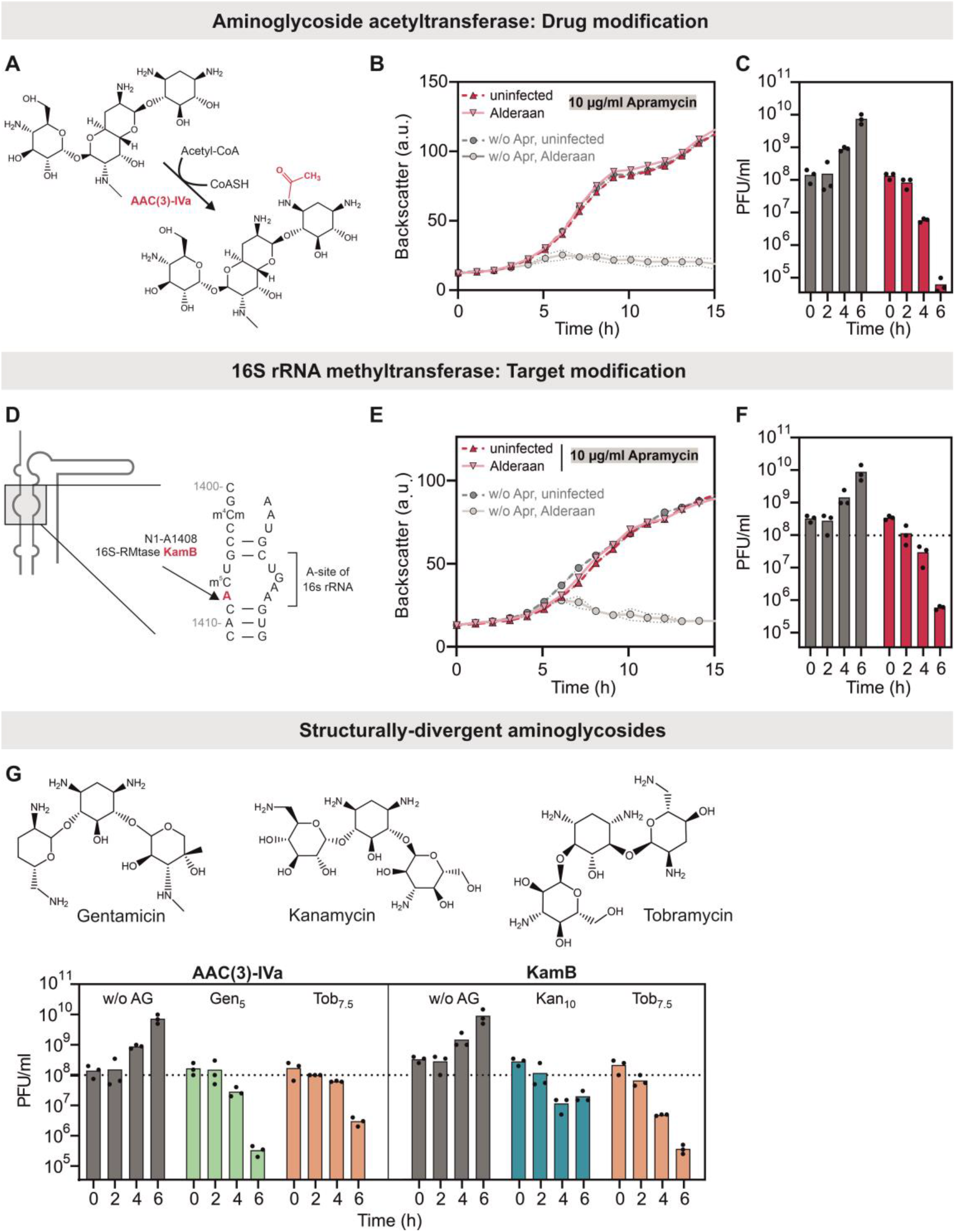
Uncoupling of antiphage and antibacterial properties of structurally-divergent aminoglycosides via drug and target site modification. A) Acetylation reaction of apramycin catalyzed by AAC(3)-IVa. B) Growth of *S. venezuelae* ATCC 10712 - pIJLK04-*aac(3)-IVa* upon infection with phage Alderaan in presence and absence of 10 µg/ml apramycin. C) Time course of phage titers during Alderaan infection shown in B). D) A-site of 16S rRNA showing the methylation position (A1408) used by methyltransferase KamB. Schematic illustration was designed according to Wachino and Arakawa (2012). E) Growth of *S. venezuelae* NRRL B-65442 – pIJ10257-*kamB* upon infection with phage Alderaan in presence and absence of 10 µg/ml apramycin. F) Time course of phage titers during Alderaan infection shown in E). G) Phage amplification during drug and target site modification via AAC(3)-IVa and KamB, respectively, in presence of gentamicin, kanamycin and tobramycin. All infection assays were conducted with an initial phage titer of 10^8^ PFU/ml in biological triplicates.

As already shown in Kever *et al*. (2022a), infection in absence of apramycin results in a complete culture collapse and a progressive phage amplification over time, whereas no more growth defect and increase in extracellular phage titer was detectable upon apramycin treatment for phage Alderaan infecting *S. venezuelae* (Figure 1B,C). A comparable extent of inhibition in terms of phage amplification and cell lysis was obtained for tobramycin and gentamicin covered by the resistance spectrum of AAC(3)-IVa as well (Figure 1G, Figure S1A-C). Although these aminoglycosides share the 2-desoxystreptamin (2-DOS) core structure with apramycin, they differ structurally in their substitutions and belong to the class of 4,6-disubstituted 2-DOS (Krause *et al*., 2016a).

To further screen for the antiphage properties of the unmodified compound, the 16S rRNA methyltransferase KamB encoded in the apramycin biosynthesis cluster of the natural producer *S. tenebrarius* was harnessed as an alternative resistance mechanism (Holmes *et al*., 1991). This methyltransferase catalyzes the N1‐methylation of the 16S rRNA at position A1408 conferring high-level resistance to the structurally-divergent aminoglycosides apramycin, kanamycin and tobramycin (Koscinski *et al*., 2007) (Figure 1D). As observed for drug modification, target site modification abolished the antibacterial mode of apramycin, but simultaneously allowed a complete inhibition of phage infection to an almost identical extent than AAC(3)-IVa (Figure 1E-F). The same applied to the other tested aminoglycosides kanamycin (4,6-disubstituted 2-DOS) and tobramycin (Figure 1G, Figure S1D-F).

For resistance via acetyltransferase AAC(3)-IVa or methyltransferase KamB, analogous results concerning the influence of aminoglycosides were also obtained for ⍰ infection of E. coli, which has previously been demonstrated to be sensitive towards apramycin and kanamycin treatment (Figure S2) (Kever *et al*., 2022a). Overall, this led us to the conclusion that acetylation neither positively nor negatively affects the antiphage activity of apramycin, gentamicin or tobramycin. Based on these findings, we infer that the inhibition of phage infection is most likely not due to a residual blockage of bacterial translation.

### Different aminoglycoside modifications do not interfere with the antiphage properties of the molecules

To examine the generality of the functional uncoupling of antiphage and antibacterial properties of aminoglycosides in more detail, we took advantage of different aminoglycoside-modifying enzymes (AMEs) targeting different positions on the aminoglycoside scaffold. In case of apramycin, just two different AMEs are described in literature including the already used acetyltransferase AAC(3)-IVa and ApmA (Figure 2**A**). ApmA reveals a unique regiospecificity by acetylating apramycin at the N2’ position of the octadiose element (Bordeleau *et al*., 2021, Bordeleau *et al*., 2023).

**Figure 2:**
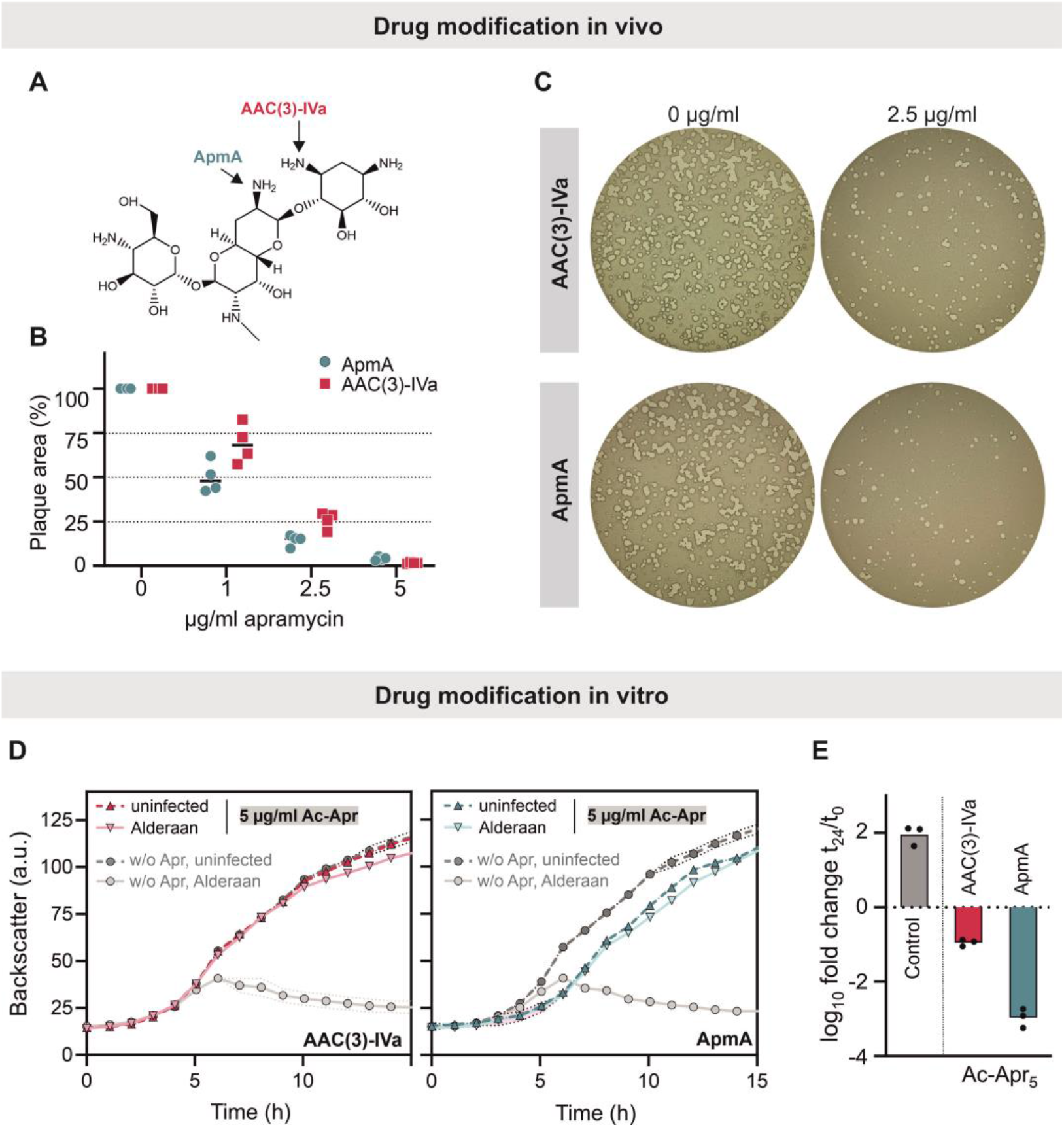
Influence of different acetylation positions on the antiphage properties of apramycin. A) Acetylation position of the two apramycin acetyltransferases AAC(3)-IVa and ApmA. B) Correlation between plaque area and apramycin concentration upon infection of the two apramycin-resistant strains *S. venezuelae* NRRL B-65442 encoding either AAC(3)-IVa or ApmA. C) Representative plaque assays to the data shown B). D) Growth of *S. venezuelae* NRRLB-65442 wildtype upon infection with phage Alderaan in presence and absence of 5 µg/ml AAC(3)-IVa- or ApmA-acetylated apramycin. E) log_10_ fold change in PFU/ml during infection shown in D). All assays were performed in biological triplicates.

Since the apramycin resistance level mediated by the acetyltransferase ApmA was reported to be more than 8-fold lower than for AAC(3)-IVa (Bordeleau *et al*., 2021), plaque assays offering higher resolution of the antiphage effect were used as a direct comparison of both modification positions. A gradual decrease in the number of plaques was detected upon increasing apramycin pressure independent of the underlying resistance gene (Figure 2B,C). This was also in line with results gained during infection of *E. coli* with phage λ. As with apramycin, different modifications of kanamycin via phosphotransferases APH(3’)-Ia and APH(2’’)-IIa or the acetyltransferase AAC(6’)-Ih did not interfere with the antiphage properties of this compound and caused a dose-dependent reduction in λ plaque formation. The same applied for modification of neomycin, an aminoglycoside with a 4,5-disubstituted 2-DOS, via APH(3’)-Ia (Figure S2).

To further verify the observed effects with a focus on apramycin effects on phage Alderaan infecting *S. venezuelae*, both acetyltransferases (ACC(3)-Iva and ApmA) were purified by affinity chromatography and used for in vitro modification of apramycin. Successful acetylation was confirmed by mass spectroscopy (Figure S3A). When supplementing acetylated apramycin to infection assays with the *S. venezuelae* wildtype strain not resistant to the antibacterial effect of apramycin, no or just a slight effect on bacterial growth could be detected for ACC(3)-IVa- and ApmA-mediated acetylation, respectively. Under infection conditions, both acetylated versions severely impacted phage infection as indicated by an omitted cell lysis and a drop in titer after 24 h of infection (Figure 2D,E), which was in line with previously published data for AAC(3)-IVa (Kever *et al*., 2022a). However, time-resolved quantification of phage titers revealed differences between the influence of intracellularly and extracellularly acetylated apramycin. In case of AAC(3)-IVa-acetylated apramycin, phage titers rose in the early stages of infection before falling below the starting level, whereas ApmA-acetylated apramycin allowed no initial increase and a more pronounced decrease in titer of ∼1000-fold after 24 h. (Figure S3B). The higher sensitivity of phage infection to ApmA-acetylated apramycin was also observed when the modified apramycin versions were added to spot assays with phage Alderaan and the *S. venezuelae* wildtype strain (Figure S3C). To conclude, together with the results presented in Figure 1, drug and target site modification were shown to efficiently uncouple the antiphage and antibacterial properties.

### Influence of spent medium of infection dynamics

Our previous study about the antiphage properties of aminoglycosides revealed that the effect of the pure compound apramycin could be reproduced with culture supernatants (=spent medium) of the natural apramycin producer *Streptoalloteichus tenebrarius* (Kever et al., 2022a). To ascertain the role of apramycin as main antiphage molecule in the supernatant interfering with phage infection and to determine the position in the biosynthetic pathway at which antiphage properties appear, we tested spent media obtained from different *S. tenebrarius* mutant strains lacking different enzymes in the biosynthetic pathway (Lv et al., 2016, Zhang et al., 2021) (Figure 3A). To analyse the impact of these spent media on phage amplification, we harnessed the apramycin-resistant *S. venezuelae* strain encoding the 16S rRNA methyltransferase KamB to prevent further modification of the apramycin biosynthesis intermediates. Resistance via target site modification allowed complete inhibition of phage infection by addition of *S. tenebrarius* wildtype spent medium (Figure 3B).

**Figure 3:**
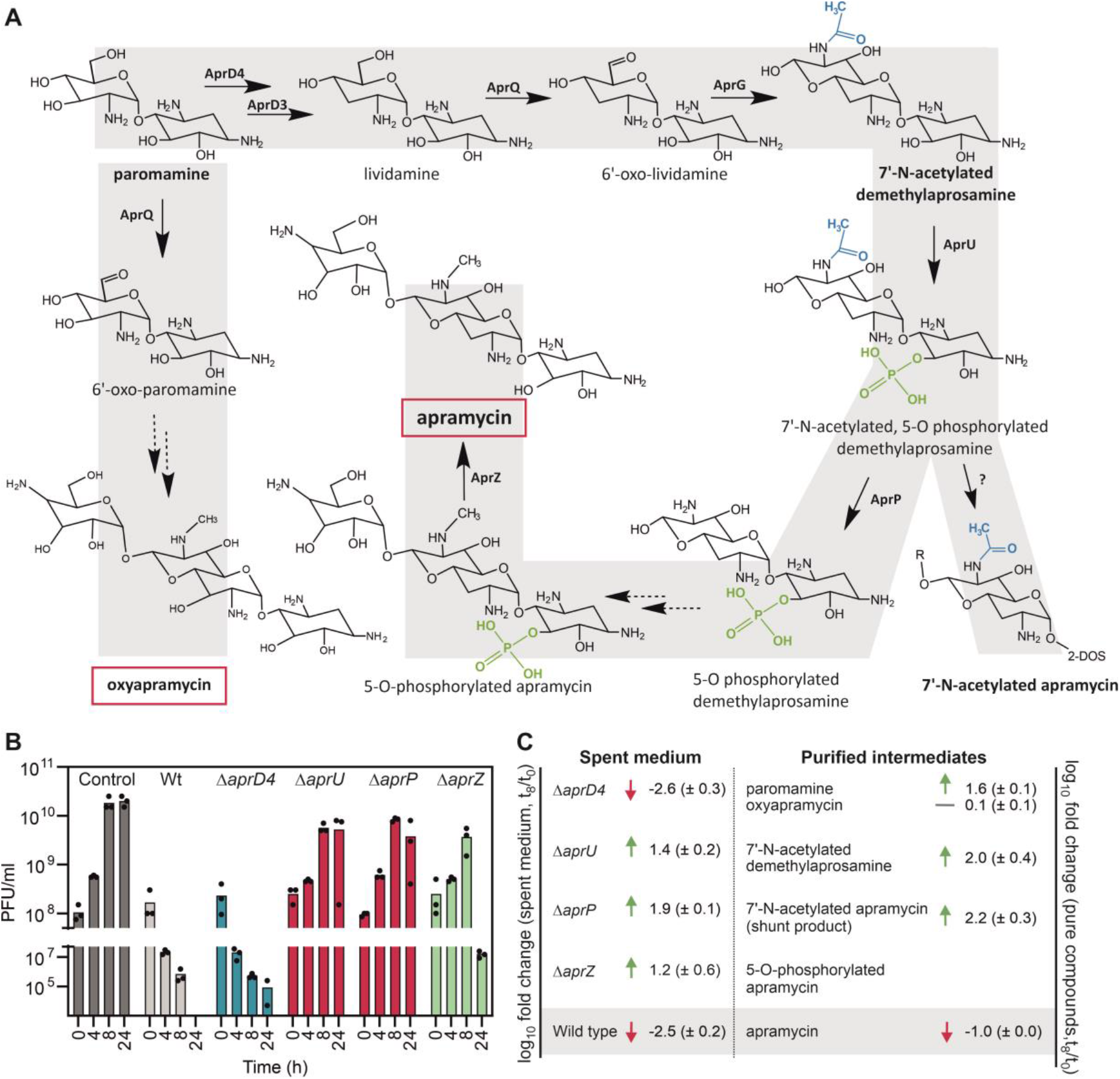
Influence of spent medium of different *S. tenebrarius* apramycin biosynthesis mutants on infection dynamics of phage Alderaan infecting *S. venezuelae* NRRL B-65442 - *kamB*. A) Biosynthetic pathway of apramycin biosynthesis in *S. tenebrarius* (2-DOS: 2.desoxystreptamin ring) (Lv *et al*., 2016, Zhang *et al*., 2021, Fan *et al*., 2023). B) Phage titers during phage infection in presence and absence of the indicated spent media. C) Log_10_ fold change (t8/t0) in PFU/ml calculated for infection in presence of spent medium and 10 µg/ml of purified biosynthesis intermediates assumed to accumulate in the different mutant strains. All assays were performed in biological triplicates.

Deletion of *aprD4* encoding a putative Fe-S oxidoreductase interferes with apramycin biosynthesis by preventing C3 deoxygenation, thereby increasing production of the apramycin analogue oxyapramycin and paromamine (Lv *et al*., 2016). Supplementing of *S. tenebrarius* Δ*aprD4* spent medium to infection assays of the apramycin-resistant *S. venezuelae* strain carrying the methyltransferase KamB showed a similar extent of phage inhibition than the wildtype spent medium. This could be traced back to the antiphage properties of oxyapramycin, as demonstrated by the addition of the purified intermediates to infection assays (Figure 3B-C, Figure S4B). Contrary to this, deletion of *aprU* encoding an aminoglycoside phosphotransferase and deletion of *aprP* encoding a putative creatinine amidohydrolase showed comparable amplification kinetics than the control infection without any supplementation, although slightly lower maximal phage titers were detected. This was in line with the successful phage infection upon addition of the acetylated demethylaprosamin and 7’-N-acetylated apramycin shown to accumulate in the *S. tenebrarius* Δ*aprU and* Δ*aprP* mutant, respectively (Figure 3B-C) (Zhang *et al*., 2021). However, comparing phage titers after 24h of infection exposed a significant reduction in plaque-forming units upon addition of acetylated apramycin reaching almost the starting level, which could not be observed for the acetylated pseudotrisaccharide or paromamine (Figure S4B). Interestingly, a similar trend could be observed for the spent medium of the Δ*aprZ* mutant lacking the extracellular alkaline phosphatase as final enzyme of the apramycin biosynthesis pathway (Zhang *et al*., 2021). Within the first 8 h, a similar extent of phage amplification was measured, but the decrease in titer after 24 h was substantially more pronounced dropping from ∼4^*^10^9^ PFU/ml at 8 h to ∼2^*^10^7^ PFU/ml at 24 h. This was accompanied by a less pronounced growth defect under infection conditions (Figure 3B-C, Figure S4A). Accordingly, intracellular formation of phosphorylated apramycin appears to prevent autotoxicity by deactivating the antibacterial properties of the compound (Zhang *et al*., 2021), but at the same time provides some degree of protection against phage infection for the producer already during apramycin biosynthesis. Subsequent extracellular dephosphorylation activates the dual functionality of apramycin, which can be hypothesized to confer community-wide protection for resistant cells in the same ecological niche. Based on the results obtained for the different mutant strains defective in apramycin biosynthesis, we conclude that the antiphage properties of apramycin and its intermediates emerge after the 7-N’-acetylated demethylaprosamine step, since not antiphage properties were observed for mutants lacking AprU or AprP.

### Co-cultivation of *S. venezuelae* and *S. tenebrarius* confirms the antiphage impact of apramycin in the context of microbial communities

To mimic community-wide antiphage defense more properly, phage-susceptible *S. venezuelae* NRRL B-65542 - *kamB* was co-cultured with apramycin-producing *S. tenebrarius*. In presence of the producer mycelium, which was pre-cultivated for two days to allow apramycin biosynthesis, Alderaan was no longer able to propagate on *S. venezuelae* (Figure 4A-B). Interestingly, phage infection could be restored by co-cultivating *S. venezuelae* with *S. tenebrarius* mycelium, which has previously been intensively washed to remove produced apramycin, while the simple addition of spent medium separated from mycelium was sufficient for phage inhibition. Furthermore, no negative effect of *S. tenebarius* mycelium and apramycin production on extracellular Alderaan phage particles was detected, further supporting the previously published hypothesis of interference with phage infection at an intracellular level (Figure 4A-B) (Kever *et al*., 2022a). In contrast, continuous phage amplification and a complete culture collapse was observed in the presence of the *S. tenebrarius* Δ*aprQ* mutant, altogether ruling out general issues in phage amplification due to the presence of other co-cultured bacteria. (Figure 4C-D). However, it is striking that no more cell growth was observed upon phage infection, even though Alderaan is not able to infect *S. tenebrarius*. This might be due to the release of as yet unidentified growth-inhibiting molecules or proteins, potentially accounting for the suppression of *S. tenebrarius* growth. In addition to showing the protective effect of *S. tenebrarius* against viral predation of *S. venezuelae*, these results highlight the need for a background level of apramycin already produced prior to phage attack.

**Figure 4:**
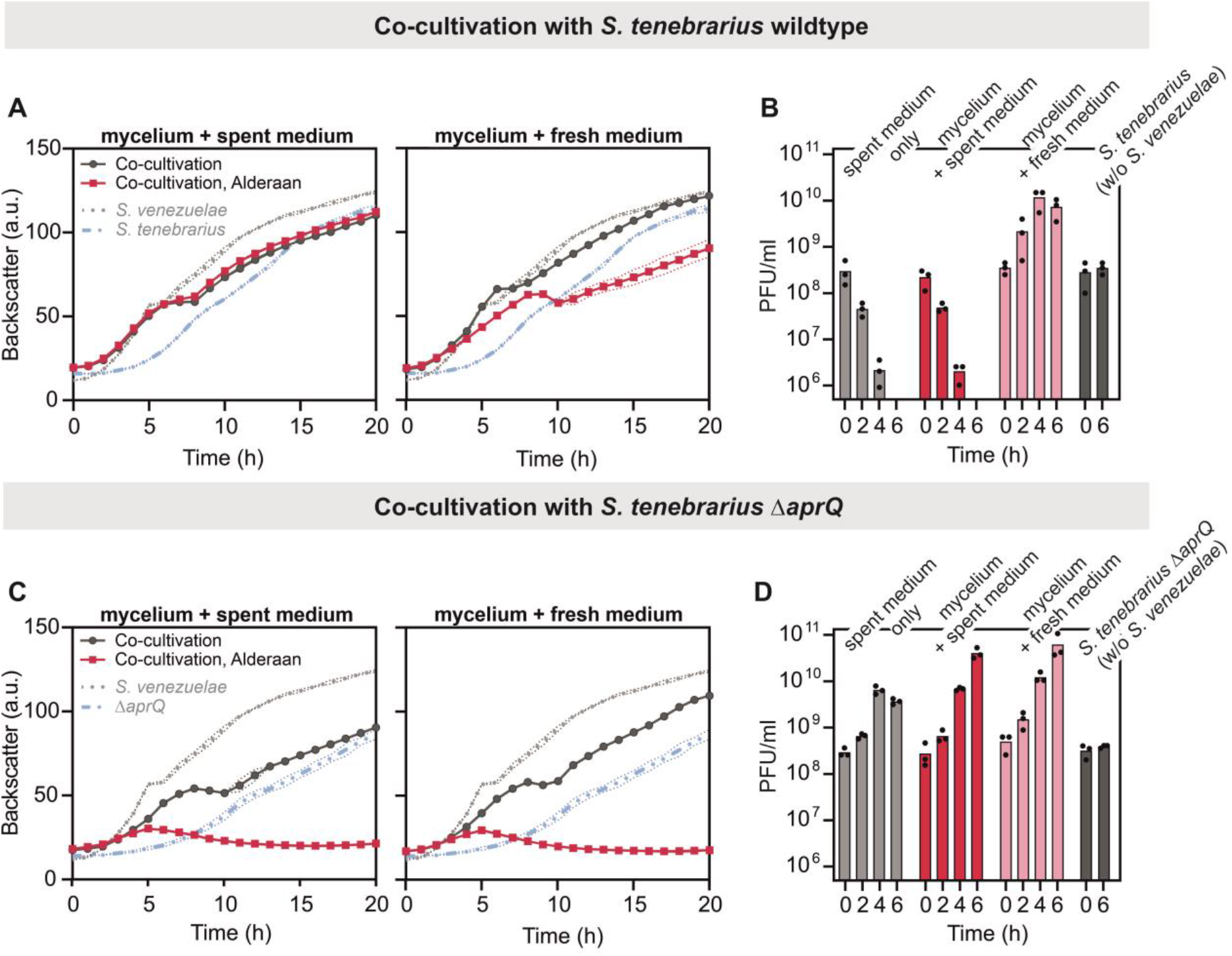
Co-cultivation of *S. venezuelae* NRRL B-65442 - *kamB* with two different *S. tenebrarius* strains during Alderaan infection. A) Infection curves during co-cultivation of *S. venezuelae* with *S. tenebrarius* wildtype directly transferred from pre-culture (+ spent medium) or after intensive washing to remove spent medium (+ fresh medium). Growth of *S. venezuelae* and *S. tenebrarius* separately from each other is shown for comparison. B) Corresponding time course of phage titers during co-cultivation and infection. Influence of *S. tenebrarius* spent medium on infection dynamics and *S. tenebrarius* itself on phage infectivity is added as control. C) Analogous data set to A) showing infection curves during co-cultivation of *S. venezuelae* with *S. tenebrarius* Δ*aprQ*. D) Corresponding time course of phage titers during co-cultivation and infection. All assays were performed in biological triplicates.

## Discussion

Aminoglycosides antibiotics were previously shown to be efficient antiphage agents in widely divergent bacterial hosts harboring an aminoglycoside-modifying enzyme as resistance mechanism (Kever *et al*., 2022a). To evaluate if such uncoupling of the dual functionality via antibacterial resistance is a more general phenomenon, we here screened a broad portfolio of aminoglycosides and resistance genes. Regardless of the use of target site or drug modification, all tested resistance mechanisms ensure bacterial growth under infection conditions by retaining the antiphage mode and abolishing the antibacterial mode of action.

As recently demonstrated by Pradier and Bedhomme (2023), aminoglycoside-resistance genes are widely distributed among all biomes, with the soil playing a pivotal role in AME gene transfer events. Natural ecosystems are considered to harbor a reservoir of antibiotic resistances genes, stemming from both, anthropogenic transfer of antibiotics and resistance genes as well as the presence of natural aminoglycoside producers such as *Streptomyces* and *Micromonospora* spp. residing in soil (Tripathi & Cytryn, 2017, Durand *et al*., 2019). Gene transfer events are facilitated by the frequent localization of such resistance genes on mobile genetic elements, allowing broadening of the resistance spectrum (Pradier & Bedhomme, 2023). Given the molecular versatility of aminoglycosides confirmed in this study, the acquisition of resistance becomes not only advantageous as defense against the chemical warfare exerted by the secretion of aminoglycosides, but could ensure concurrent protection from phages as already discussed in Hardy *et al*. (2023). The prevalence of aminoglycoside-resistant bacteria in natural habitats puts further emphasis on the ecological significance of such chemical antiphage defense. However, taken into account the predominance of sub-inhibitory antibiotic concentrations and the existence of local extremes in the soil environment (Fajardo *et al*., 2009), one can infer that such community-wide antiphage defense is constrained to mutualistic interactions within the genetic kin or between different species. The latter is thought to be facilitated by increased horizontal gene transfer events of aminoglycoside resistance genes in the presence of low antibiotic concentrations (Cairns *et al*., 2018). Overall, this would add to the notion of the pan-immune system describing antiphage defense as a shared community resource (Bernheim & Sorek, 2020).

An additional aspect, which reinforces the notion of aminoglycosides as “public goods” in antiphage defense is the natural growth phenotype as well as the temporal onset and organization of antibiotic production in *Streptomyces* spp. These multicellular growing microorganisms are characterized by a complex developmental program, starting with the germination of a spore and the establishment of a vegetative mycelium, which further develop into aerial hyphae and spores upon nutrient depletion (Bush *et al*., 2015, Schlimpert & Elliot, 2023). Antibiotic production is typically linked to the morphological differentiation to aerial mycelium in order to protect released nutrients of sacrified mycelial parts from other terrestrial competitors (van der Meij *et al*., 2017, Bibb, 2005). To enhance colony-wide fitness, this antibiotic production can be coordinated by a division of labor, involving the differentiation into an altruistic subpopulation specialized in antibiotic production (Zhang *et al*., 2020, Zhang *et al*., 2022), which in turn is facilitated by the genomic instability of *Streptomyces* at the chromosomal arms (Thibessard & Leblond, 2014, Bentley *et al*., 2002). Given the clonal nature of the *Streptomyces* colonies, secretion of aminoglycosides might play a pivotal role in shielding susceptible segments of the colony against both bacterial competitors and phage predation.

In addition to the ecological relevance, this study provides structural insights into the antiphage effect of these compounds. According to assays with the different *S. tenebrarius* mutant strains and purified apramycin biosynthetic intermediates, the antiphage effect occurred at a late step of the biosynthetic pathway (Figure 3). This could be due to either differences in the intracellular interaction with phage amplification or in the uptake of the respective molecules. Modification via acetylation or phosphorylation reduces the amount of positively charged amino groups at neutral pH or even adds a negative charge to the cationic aminoglycosides, which could affect their interaction with the negatively charged components of the bacterial surface for uptake (Taber *et al*., 1987). In the case of the in vitro-acetylated apramycin, the interference with phage infection was less pronounced in the early stages of infection, but comparable to the in vivo modification in the long-term, supporting the hypothesis of a reduced uptake of the molecules when modified in vitro. The same could apply to the phosphorylated apramycin accumulating in the Δ*aprZ* mutant, possibly explaining the delayed phage-inhibiting effect. These considerations highlight the importance of discriminating between differences in molecule uptake and a potential antiphage property exerted at the intracellular level (Kever *et al*., 2022b). Aminoglycosides are known to bind to the bacterial cell membrane and subsequently accumulate within the cell via electron-transport mediated processes, EDPI and EDPII (Krause *et al*., 2016b). Using fluorescently labelled antibiotics, the membrane-bound and EDPII elements were shown to constitute a substantial proportion of the overall fluorescence levels tightly bound to *E. coli* cells (Sabeti Azad *et al*., 2020). In cells expressing resistance genes, one could reasonably expect to predominantly encounter the membrane-bound fraction of aminoglycosides. This anticipation arises from the assumption that the disruption of membrane integrity and the diffusive entry of the drug, induced by mistranslated proteins during EDPII, would not be anticipated in the presence of resistance mechanisms.

While the acquisition of aminoglycoside resistance holds several advantages for bacteria, it poses a drawback for antibiotic-phage combinatory treatments (Jiang *et al*., 2020, Zuo *et al*., 2021). Although the clinical use of aminoglycosides declined with the introduction of other classes of antibiotics such as cephalosporins or fluoroquinolones, the global spread of multidrug-resistant pathogens has led to a renewed demand for this class of antibiotics, frequently for combinatory treatments (Krause *et al*., 2016a). Moreover, increasing knowledge about modifications overcoming aminoglycoside-resistance allows design of semi-synthetic antibiotics, thereby improving the therapeutic window of aminoglycosides to combat this global health issue (Serio *et al*., 2018, Zárate *et al*., 2018, Krause *et al*., 2016a). However, attention should be paid to the pre-selection of phages for antibiotic-phage combinatory treatments to avoid antagonistic effects, since uncoupling of antiphage and antibacterial effects seems feasible with divergent aminoglycoside resistance genes.

In conclusion, this study emphasizes the ecological relevance of natural aminoglycoside secretion for a community-wide defense against phages by highlighting the functional decoupling via aminoglycoside-resistance genes. Moreover, it sheds light on crucial considerations for the integration of phage-antibiotic combinatory treatments in the context of clinical infections.

## Material and methods

### Bacterial strains and growth conditions

All resistance cassettes, bacterial strains and phages used in this study are listed in Table S1 and S2. For *Streptomyces venezuelae*, pre-cultures for growth and infection assays were inoculated from spore stocks in glucose-yeast extract-malt extract (GYM) medium containing 50% tap water, pH 7.3 and cultivated at 30°C and 120 rpm for 18 h. Pre-cultures were subsequently used to inoculate main cultures in the same medium to the indicated OD_450_. For *E. coli*, cultures were inoculated from a single clone grown on agar plates and cultivated at 170 rpm and 37 °C O/N in LB medium for being subsequently used to inoculate main cultures to the indicated OD_600_. In all cases, antibiotics were added to cultures as indicated.

For standard cloning procedures and protein overproduction *E. coli* DH5α and *E. coli* BL21 (DE3) were used, respectively. Infection assays were conducted in *E. coli* LE392 and conjugation between *Streptomyces* spp. and *E. coli* was done via the conjugative *E. coli* ET12567/pUZ8002.

In order to determine the titer of plaque-forming units during infection assays, spot assays were conducted. To this end, 2 µl of decimal dilution series of culture in sodium chloride/magnesium sulfate (SM) buffer (10 mM Tris-HCl pH 7.3, 100 mM NaCl, 10 mM MgCl_2_, 2 mM CaCl_2_) were spotted on a bacterial lawn propagated on a double-agar overlay with the top layer inoculated to an optical density of OD_450_ = 0.3 and OD_600_ = 0.2 for *S. venezuelae* and *E. coli*, respectively. Plaque assays with *S. venezuelae* were done in a comparable manner by adding 10^3^ PFU/ml of phage Alderaan to GYM soft-agar inoculated with mycelium to an OD_450_ of 0.3. Plaque areas were analysed using ImageJ and the color thresholding tool. To this end, images were cropped and the color threshold settings were adjusted to select plaques as accurately as possible. Subsequently, plaque areas were measured using the “analyze particle” option with the following parameters: size (pixel^2^): 40 to infinity, circularity: 0.1-1.0.

### Recombinant DNA work and cloning

All plasmids and oligonucleotides used in this study are listed in Table S3 and S4. Standard cloning techniques such as PCR, restriction digestion with indicated restrictions enzymes and Gibson assembly were performed according to standard protocols **(Sambrook & Russell, 2001, Gibson, 2011)**. DNA sequencing as well as synthesis of oligonucleotides and (codon-optimized) genes was performed by Eurofins Genomics (Ebersberg, Germany).

### Phage infection curves

Growth assays and phage infection curves were performed in the BioLector microcultivation system (Beckman Coulter Life Sciences (formerly m2p-labs), Krefeld, Germany) using a shaking frequency of 1200 rpm and cultivation temperature of 30°C. Biomass was measured as a function of backscattered light intensity with an excitation wavelength of 620 nm (filter module: λ_Ex_/ λ_Em_: 620 nm/ 620 nm, gain: 25) from three independent biological replicates.

Cultivation of *S. venezuelae* was performed in 1 ml GYM medium (pH 7.3, 50% tap water) inoculated with an overnight culture to a starting OD_450_ of 0.15. Infection was conducted by supplementing the indicated initial phage titers to the respective wells. Culture supernatants were collected in specific time intervals to monitor phage amplification over time via double-agar overlay assays.

### Co-cultivation in submerged cultures

*S. tenebrarius* strains were inoculated from mycelial stocks in 5 ml GYM medium (pH 7.3, 50% tap water) and cultivated at 170 rpm and 30 °C for 24 h. Subsequently, these pre-cultures were used to inoculate 20 ml GYM medium (pH 7.3, 50% tap water) to an initial OD_450_ of 0.2. Cultures were incubated for 3 days at 120 rpm and 30 °C to allow production and secretion of apramycin or its biosynthesis intermediates. One day prior to the co-cultivation assay, *S. venezuelae* NRRL B-65442-*kamB* was inoculated from spores in 5 ml GYM medium (pH 7.3, 50% tap water) and cultivated at 170 rpm and 30 °C for 24 h. For co-cultivation, 1 OD unit of *S. tenebrarius* mycelium was mixed with 1 OD unit of *S. venezuelae* mycelium in 1 ml GYM medium (pH 7.3, 50% tap water). Cultivation was performed in the BioLector microcultivation system as described in ‘Phage infection curves’ in biological triplicates. Phage infection was performed by addition of 10^7^ PFU/ml Alderaan particles. As comparison, mycelium of *S. tenebrarius* was either directly transferred to co-cultivations or washed twice beforehand to remove produced metabolites.

### Preparation of culture supernatants (referred to as spent media)

In order to collect spent medium of the natural apramycin producer *Streptoalloteichus tenebrarius* and respective biosynthesis mutants, all strains were cultivated as described in ‘Co-cultivation in submerged cultures’ using a final cultivation volume of 70 ml GYM medium (pH 7.3, 50% tap water). Culture supernatants were harvested by centrifugation at 5,000 g at 4 °C for 20 min and subsequently sterile filtered before storage at 4°C for direct usage in infection assays.

### Infection assays in spent medium

Infection assays of apramycin-resistant *S. venezuelae* in spent medium were conducted in the BioLector microcultivation system (Beckman Coulter Life Sciences (formerly m2p-labs), Krefeld, Germany) as described in ‘Phage infection curves’ with the following modifications: 1.25x GYM medium (pH 7.3, 50% tap water) was supplemented with spent medium to a final concentration of 20% (v/v).

### Purification of biosynthesis intermediates

Paromamine and oxyapramycin were generated from Δ*aprD4* mutant WDY288 (Lv *et al*., 2016). Acetylated apramycin and acetylated pseudotetrasaccharide were generated from Δ*aprP* mutant WDY321 (Zhang *et al*., 2021). The two mutant strains were cultivated on SPA medium (2% soluble starch, 0.1% beef extract, 0.05% MgSO_4_, 0.1% KNO_3_, 0.05% NaCl, 0.05% K_2_HPO_4_, 2% agar) at 37 °C for spore production. Their seed culture was prepared in 5 ml TSBY medium (3% tryptone soya broth and 0.5% yeast extract) at 37 °C with shaking at 220 rpm for 2 days. The seed culture of Δ*aprD4* and Δ*aprP* were sub-cultured into 50 mL SPC fermentation medium (4% glucose, 1% peptone, 0.4% soybean meal, 1% corn meal, 0.4% MgSO_4_^*^7H_2_O, 0.5% NH4Cl, 0.05% FeSO4, 0.03% MnCl2, 0.003% ZnSO_4_,0.5% CaCO_3_) at 37°C with shaking at 220 rpm for 7 days. The culture supernatant was collected by centrifugation at 5000 rpm for 30 min. Afterwards, the supernatant was adjusted to a pH of 2-3 with saturated oxalate and the insoluble fraction was removed by centrifugation at 5,000 rpm for 30 min. Next, the supernatant was loaded onto the 732 cation exchange resin (Hebi Juxing Resinco., Ltd, Hebi, China), before being eluted by 3% ammonia hydroxide solution. Finally, the ammonia hydroxide eluate was purged with a nitrogen stream.

The filtered eluate was concentrated and subjected to a semi-preparative HPLC System equipped with Evaporative Light Scattering Detector (ELSD, Alltech 2000ES) and an Agilent ZORBAX SB-C18 column (5 μm, 250×9.4mm). Gradient elution was performed at a flow rate of 3.0 ml/min. The gas flow and temperature of ELSD was set to 2.9 l/min and 109 °C. Gradient elution was performed at a flow rate of 3 ml/min with solvent A (water containing 10 mM heptafluorobutyric acid) and solvent B (CH_3_CN): 0-3 min, constant 80% A/20% B; 3-5 min, a linear gradient to 75% A/25% B; 5-9 min, a linear gradient to 70% A/30% B; 9-17 min, a linear gradient to 59% A/41% B; 17-18 min, a linear gradient to 80% A/20% B; 18-25 min, constant with 80% A/20% B. The eluent containing the target components was collected and subsequently dried through lyophilization.

### Protein purification via affinity chromatography

For heterologous protein overproduction, *E. coli* BL21 (DE3) cells containing Strep-tagged versions of the respective resistance genes on the pAN6 plasmid were pre-cultivated in LB medium supplemented with 50 µg/ml kanamycin (LB Kan_50_) at 37 °C and 120 rpm. The pre-culture was used to inoculate the main culture in the same medium to an OD_600_ of 0.1 and cultivation was continued at 37°C, and 120 rpm until gene expression was induced with 100 μM IPTG at an OD_600_ of 0.6. Cells were harvested after further cultivation at 120 rpm and 18 °C for 16 h.

Cell harvesting and disruption were performed with the multi shot cell disruptor at 20,000 psi using buffer A (100 mM Tris-HCl, pH 8.0) with cOmplete™ Protease inhibitor (Roche, Basel, Switzerland) for cell resuspension and disruption. After centrifugation at 20,000 g and 4 °C for 30 min buffer B (100 mM Tris-HCl, 500 mM NaCl, pH 8.0) was used for purification.

To this end, supernatant containing either the C-terminal Strep-tagged ApmA or the AAC(3)-IVa apramycin acetyltransferase was applied to an equilibrated 2 ml Strep-Tactin-Sepharose column (IBA, Göttingen, Germany). After washing with 30 ml buffer B, the protein was eluted with 5 ml buffer B containing 15 mM D-desthiobiotin (Sigma Aldrich, St. Louis, USA). After purification, elution fractions with the highest protein concentrations were pooled and checked by SDS-PAGE (Laemmli, 1970) using a 4–20% Mini-PROTEAN gradient gel (BioRad, Munich, Germany). Final protein concentration of the pooled elution fraction was determined with the Pierce BCA Protein Assay Kit (ThermoFisher Scientific, Waltham, MA, USA) before being used for in vitro acetylation.

### In vitro acetylation of apramycin

Purification of the two acetyltransferases AAC(3)-IVa and ApmA was performed as described above. Acetylation of apramycin was done as described in Kever *et al*. (2022a) with the following modifications. Assay mixtures were composed of 350 µl 100 mM Tris-HCl (pH 8.0), 50 µl purified enzyme (end concentration: 30 µg/mL), 50 µl apramycin (end concentration: 10 mM) and 50 µl acetyl-CoA trisodium salt (SigmaAldrich, St. Louis, MO, USA; end concentration: 10 mM). The assay mixtures were incubated at 37°C for 30 min.

### Direct injection mass spectrometry

For checking acetylation efficiency, direct injection mass spectrometry was performed. To this end, samples were diluted 1:100 (v/v) with deionized water and further 1:10 (v/v) with 50% methanol containing 0.1% acetic acid. The individual samples were directly injected into the HESI-source of a QExactive Plus mass spectrometer (Thermo Fisher) using a 500µl syringe with a constant flow of 5 µl/min. The source parameters were set as follows: spray voltage (+): 3200 V, capillary temperature: 320.00 °C, sheath gas: 8.0, aux gas: 2.0, S-Lens RF Level: 50.00. Once the spray stabilized data were acquired for 2 min in positive mode with a full scan resolution of 70,000 and an AGC target of 3e6. Representative spectra data were retrieved from mid run scans for each sample.

## Author Contributions

Conceptualization: LK, AH, JF; Data Curation: LK; Formal Analysis: LK, QZ, PW; Funding acquisition: YY, JF; Investigation: LK, QZ, AH, PW; Methodology: All; Project administration: JF; Resources: YY, JF; Supervision: YY, JF; Validation: All; Visualization: LK; Writing—original draft: LK; Writing—review and editing: All; All authors have read and agreed to the published version of the manuscript.

## Acknowledgements

We thank the Deutsche Forschungsgemeinschaft (SPP 2330, project 464434020 and SFB 1535, project ID 458090666) for financial support.

