## Supplementary material for "Decoupling the molecular versatility of aminoglycosides via drug or target modification enables community-wide antiphage defense": Tables S1-4 and Figures S1-4

Table S1: Aminoglycoside resistance mechanisms used in this study

Table S2: Bacterial strains and phages

Table S3: Plasmids used in this study

Table S4: Oligonucleotides used in this study

**Figures**

Figure S1: Inhibition of phage Alderaan infection by structurally-divergent aminoglycosides in aminoglycoside-resistant *S. venezuelae* strains.

Figure S2: Influence of different aminoglycoside modifications on phage $\lambda$ infection.

Figure S3: Acetylation of apramycin with the two purified acetyltransferases AAC(3)-IVa and ApmA and its influence on phage infection dynamics.

Figure S4: Growth curves upon infection of *S. venezuelae* NRRL B-65442 - *kamB* with phage Alderaan in presence of spent medium from different *S. tenebrarius* strains.

**Table S1: Aminoglycoside resistance mechanisms used in this study.**

| Resistance gene | Host organisms | Protein sequence | Resistance profile |
|---|---|---|---|
| APH(3')-Ia | *Escherichia coli, Klebsiella pneumoniae* | MSHIQRETSCSRPRLNSNMDADLYGYKW ARDNVGQSGATIYRLYGKPDAPELFLKHG KGSVANDVTDEMVRLNWLTEFMPLPTIK HFIRTPDDAWLLTTAIPGKTAFQVLEEYPD SGENIVDALAVFLRRLHSIPVCNCPFNSDR VFRLAQAQSRMNNGLVDASDFDDERNG WPVEQVWKEMHKLLPFSPDSVVTHGDF SLDNLIFDEGKLIGCIDVGRVGIADRYQDL AILWNCLGEFSPSLQKRLFQKYGIDNPDM NKLQFHLMLDEFF* | Kanamycin, Neomycin |
| AAAC(6')-Ih | *Actinobacter baumanii* | MNIMPISESQLSDWLALRCLLWPDHEDV HLQEMRQLITQAHRLQLLAYTDTQQAIA MLEASIRYEYVNGTQTSPVAFLEGIFVLPEY RRSGIATGLVQQVEIWAKQFACTEFASDA ALDNQISHAMHQALGFHETERVVYFKKNI G* | Kanamycin, Tobramycin |
| APH(2'')-Iia | *Campylobacter jejuni* | MIDLDVEIYQHLNEQIKINKLCYLSSGDDS DTFLCNEQYVVKVPKRDSVRFAQKREFEL YRFLENCNLSYQTPAVVYQSDRFNIMKYIK GERITYEQYHKLSEKEKDALAYDEATFLKEL HSIEIDCSVSLFSDALVNKKDKFLQDKKLLIS ILEKEQLLTDEMLEHIETIYENISSNAVIFNYI PCLVHNDFSANNLIFRNNRLFGVIDFGDF NVGDPDNDFLCLLDCSTDDFGKEFGRKVL KYYQHKAPEVAERKAELNDVYWSIDQIIY GYERKDREMLIKGVSELLQTQAEMFIF* | Kanamycin, Tobramycin |
| AAC(3)-IVa | *Salmonelle enterica* | VQYEWRKAELIGQLLNLGVTPGGVLLVHS SFRSVRPLEDGPLGLIEALRAALGPGGTLV MPSWSGLDDEPFDPATSPVTPDLGVVSD TFWRLPNVKRSAHPFAFAAAGPQAEQIIS DPLPLPPHSPASPVARVHELDGQVLLLGV GHDANTTLHLAELMAKVPYGVPRHCTILQ DGKLVRVDYLENDHCCERFALADRWLKE KSLQKEGPVGHAFARLIRSRDIVATALGQL GRDPLIFLHPPEAGCEECDAARQSIG* | Apramycin, Tobramycin, Gentamicin |
| ApmA | *Staphylococcus aureus* | MKTRLEQVLERYLNGREVAVWGVPTRRL LRALKPFKFHTADRVDPQYHYVVAVTDD DLTDFLSDEQSKSFQYANDYLTFDDEGGE LPFERMCFNVPVGRQTYFGDGVVGACEN GYIKSIGQFTSINGTAEIHANHQLNMTFVS DDIQNFFNEESMAVFQEKLRKDPKHPYAY SKEPMTIGSDVYIGAHAFINASTVTSIGDG AIIGSGAVVLENVPPFAVVVGVPARIKRYR FSKEMIETLLRVKWWDWSIEEINENVDAL ISPELFMKKYGSL* | Apramycin |

| | | | |
|---|---|---|---|
| KamB | *Streptoalloteichus tenebrarius* | MRRVVGKRVQEFSDAEFEQLRSQYDDVVLDVGTGDGKHPYKVARQNPSRLVVALDADKSRMEKISAKAAAKPAKGGLPNLLYLWATAERLPPLSGVGELHVLMPWGSLLRGVLGSSPEMLRGMAAVCRPGASFLVALNLHAWRPSVPEVGEHPEPTPDSADEWLAPRYAEAGWKLADCRYLEPEEVAGLETSWTRRLHSSRDRFDVLALTGTISP* | Apramycin, Kanamycin, Tobramycin |

**Table S2: Bacterial strains and phages**

| Strain | Characteristics | Reference |
|---|---|---|
| ***Streptomyces venezuelae* ATCC 10712** | Wildtype strain | (Ehrlich et al., 1948) |
| ***S. venezuelae* ATCC 10712 – *aac(3)IV*** | *S. venezuelae* ATCC 10712 carrying the integrative plasmid pIJLK04-*aac(3)IV*, Apr$^{R}$ | (Kever et al., 2022) |
| ***Streptomyces venezuelae* NRRL B-65442** | Wild type strain | (Gomez-Escribano et al., 2021) |
| ***S. venezuelae* NRRL B-65442 – *aac(3)IV*** | *S. venezuelae* NRRL B-65442 carrying the integrative plasmid pIJLK04-*aac(3)IV*, Apr$^{R}$ | This study |
| ***S. venezuelae* NRRL B-65442 – *apmA*** | *S. venezuelae* NRRL B-65442 carrying the integrative plasmid pIJLK06-*apmA*, Apr$^{R}$ | This study |
| ***S. venezuelae* NRRL B-65442 – *kamB*** | *S. venezuelae* NRRL B-65442 carrying the integrative plasmid pIJ10257-*kamB*, Apr$^{R}$ | This study |
| ***Escherichia coli* LE392** | *$F^-$ hsdR514 ($rk^-$ $mk^-$) supE44 supF58 Δ(lacIZY)6 galK2 galT22 metB1 trpR55 $lambda^-$* | (Murray et al., 1977) |
| ***E. coli* LE392 – *aph(3')-Ia*** | *E. coli* LE392 carrying the plasmid pEKEx2a-*aph(3')-Ia*, Kan$^{R}$ | (Kever et al., 2022) |
| ***E. coli* LE392– *aac(3)IV*** | *E. coli* LE392 carrying the plasmid pEKEx2d-*aac(3)IV*, Apr$^{R}$ | (Kever et al., 2022) |
| ***E. coli* - *kamB*** | *E. coli* LE392 carrying the plasmid pEKEx2f-*kamB*, Kan$^{R}$ | This study |
| ***E. coli* LE392 - *aac(6')-Ih*** | *E. coli* LE392 carrying the plasmid pEKEx2i-*aac(6')-Ih*, Kan$^{R}$ | This study |
| ***E. coli* - *aph(2'')-IIa*** | *E. coli* LE392 carrying the plasmid pEKEx2l-*aph(2'')-Ib*, Kan$^{R}$ | This study |
| ***E. coli* LE392– *apmA*** | *E. coli* LE392 carrying the plasmid pEKEx2m-*apmA*, Apr$^{R}$ | This study |
| ***E. coli* ET12567 pUZ8002** | *dam-13::Tn9 dcm-6 hsdM hsdR,* carrying plasmid pUZ8002 | (MacNeil et al., 1992) |
| ***Streptoalloteichus tenebrarius* ATCC 17920** | Wildtype strain | (Higgins & Kastner, 1967) |
| ***S. tenebrarius* aprD4** | In-frame deletion of the *aprD4* gene | (Lv et al., 2016) |
| ***S. tenebrarius* aprP** | In-frame deletion of the *aprP* gene | (Zhang et al., 2021) |
| ***S. tenebrarius* aprQ** | In-frame deletion of the *aprQ* gene | (Lv et al., 2016) |
| ***S. tenebrarius* aprU** | In-frame deletion of the *aprU* gene | (Zhang et al., 2021) |
| ***S. tenebrarius* aprZ** | In-frame deletion of the *aprZ* gene | (Zhang et al., 2021) |
| **Alderaan** | Virulent phage infecting *S. venezuelae*, dsDNA | (Hardy et al., 2020) |
| **Lambda** | Temperate phage infecting *E. coli,* dsDNA | DSM4499 |

**Table S3: Plasmids used in this study.**

| Plasmid | Characteristics | Reference |
|---|---|---|
| **pIJ10257** | $Hyg^R$; Cloning vector for the conjugal transfer of DNA from *E. coli* to *Streptomyces spp.;* (constitutive promoter *ermE**; Integration at the ΦBT1 attachment site) | (Hong et al., 2005) |
| **pIJLK01** | $Hyg^R$; Derivative of pIJ10257 with additional restrictions sites Bst1107I (upstream) and StuI (downstream) of the *aph(7'')-Ia* gene allowing exchanging of the antibiotic cassette | (Kever et al., 2022) |
| **pIJLK04 – *aac(3)IV*** | $Apr^R$; Derivative of pIJLK01 with *aph(7'')-Ia* exchanged for *aac(3)IV* (apramycin resistance gene) | (Kever et al., 2022) |
| **pEKEx2** | $Kan^R$; *C. glutamicum*/ *E. coli* shuttle vector for regulated gene expression; $P_{tac}$, *lacI*$^q$, pBL1 oriV$_{C.g.}$, pUC18 oriV$_{E.c.}$ | (Eikmanns et al., 1991) |
| **pEKEx2a – *aph(3')-Ia*** | $Kan^R$; Derivative of pEKEx2 with additional restrictions sites Bst1107I (upstream) and NotI (downstream) of the *aphA1* gene allowing exchanging of the antibiotic cassette | (Kever et al., 2022) |
| **pEKEx2d – *aac(3)IV*** | $Apr^R$; Derivative of pEKEx2a with *aphA1* exchanged for *aac(3)IV* (apramycin resistance gene) | (Kever et al., 2022) |
| **pAN6** | $Kan^R$.; *E. coli* vector for regulated gene expression; derivative of pEKEx2 ($P_{tac}$, *lacI*$^q$, pBL1 *oriV*$_{C.g.}$, pUC18 *oriV*$_{E.c.}$) | (Frunzke et al., 2008) |
| **pAN6-TEV-Cstrep** | $Kan^R$.; Derivative of pAN6 with TEV cleavage site and C-terminal Strep-tag | Davoudi, unpublished |
| **pAN6-*aac(3)IV*-Cstrep** | $Kan^R$; Derivative of pAN6 with *aac(3)IV* fused to a C-terminal Strep-tag | (Kever et al., 2022) |
| **pUZ8002** | $Kan^R$; RK2 derivative with nontransmissible oriT | (Paget et al., 1999) |

| Plasmid | Characteristics | Template | Primer | Vector | Restriction enzyme | Sequencing primer |
|---|---|---|---|---|---|---|
| pIJLK06 – *apmA* | $Apr^R$; Derivative of pIJLK01 with *aph(7'')-Ia* exchanged for *apmA* (apramycin resistance gene) | Synthesized *apmA* gene | 1 + 2 | pIJLK01 | Bst1107I; StuI | 15 + 16 |
| pIJ10257 – *kamB* | $Apr^R$; Derivative of pIJ10257 with *kamB under the control of the constitutive promotor ermE** (apramycin resistance gene | gDNA of *S tenebrarius* | 3 + 4 | pIJ10257 | NdeI; HindIII | 17 + 18 |
| pEKEx2f – *kamB* | $Apr^R$; Derivative of pEKEx2a with *aph(3')-Ia* exchanged for *kamB* (apramycin resistance gene) | Synthesized *kamB* gene, codon-optimized | 5 + 6 | pEKEx2a | Bst1107I; NotI | 19 + 20 |

| | | | | | | |
|---|---|---|---|---|---|---|
| pEKEx2i – *aac(6')-Ih* | Apr$^R$; Derivative of pEKEx2a with *aph(3')-Ia* exchanged for *aac(6')-Ih* (kanamycin resistance gene) | Synthesized *aac(6')-Ih* gene | 7 + 8 | pEKEx2a | Bst1107I; NotI | 19 + 20 |
| pEKEx2l – *aph(2'')-IIa* | Apr$^R$; Derivative of pEKEx2a with *aph(3')-Ia* exchanged for *aph(2'')-IIa* (kanamycin resistance gene) | Synthesized *aph(2'')-IIa* gene, codon-optimized | 9 + 10 | pEKEx2a | Bst1107I; NotI | 19 + 20 |
| pEKEx2m – *apmA* | Apr$^R$; Derivative of pEKEx2a with *aph(3')-Ia* exchanged for *apmA* (apramycin resistance gene) | Synthesized *apmA* gene, codon-optimized | 11 + 12 | pEKEx2a | Bst1107I; NotI | 19 + 20 |
| pAN6 – *apmA*-Cstrep | Kan$^R$; Derivative of pAN6 with *apmA* fused to a C-terminal Strep-tag | Synthesized *apmA* gene, codon-optimized | 13 + 14 | pAN6-TEV-Cstrep | NheI; NdeI | 21 + 22 |

**Table S4: Oligonucleotides used in this study.**

| No. | Name | Oligonucleotide sequence (5’-3’) |
|---|---|---|
| 1 | **pIJLK06 – *apmA* - fw** | GTGAATAGAGGTCCGCTGTATACATGAAAACCAGACTTGAACAAGTTTTAGAAC |
| 2 | **pIJLK06 – *apmA* - rv** | CCGGGCGGCCCGGGGCGAGGCCTTTACAAACTCCCGTACTTTTTCATAAATAGTTCAG GTG |
| 3 | **pIJ10257 – *kamB - fw*** | TAGAACAGGAGGCCCCATATGATGCGCCGCGTGGTGGGCAA |
| 4 | **pIJ10257 – *kamB - rv*** | TCATGAGAACCTAGGATCCAAGCTTTCACGGACTGATCGTGCCGGTGAG |
| 5 | **pEKEx2f – *kamB* - fw** | GTAATACAAGGGGTGTTGTATACATGCGTCGTGTGGTTGGCAA |
| 6 | **pEKEx2f – *kamB* - rv** | AATTAACCAATTCTGAGCGGCCGCTTACGGTGAAATGGTGCCAGTCA |
| 7 | **pEKEx2i – *aac(6’)-Ih* -fw** | TACAAGGGGTGTTGTATACATGAATATTATGCCGATATCTGAATCACAA |
| 8 | **pEKEx2i – *aac(6’)-Ih* - rv** | ACCAATTCTGAGCGGCCGCTTAGCCGATATTTTTCTTAAAATACACCAC |
| 9 | **pEKEx2l – *aph(2’’)-IIa* - fw** | GTAATACAAGGGGTGTTGTATACATGATCGATCTGGACGTGGAAATCT |
| 10 | **pEKEx2l – *aph(2’’)-IIa* - rv** | TTAACCAATTCTGAGCGGCCGCTTAAAAGATGAACATTTCAGCTTGTGT |
| 11 | **pEKEx2m – *apmA* - fw** | GTAATACAAGGGGTGTTGTATACATGAAAACACGCCTGGAACAA |
| 12 | **pEKEx2m – *apmA* - rv** | AATTAACCAATTCTGAGCGGCCGCTTAGAGACTCCCGTATTTCTTCATG |
| 13 | **pAN6 – *apmA- Cstrep* - fw** | ATGCCTGCAGAAGGAGATATACATATGATGAAAACACGCCTGGAACAAG |
| 14 | **pAN6 – *apmA- Cstrep* - rv** | TGAAAATACAGGTTCTCGCTAGCGAGACTCCCGTATTTCTTCATGAACA |
| 15 | **pIJLK – seq – fw** | CGTAGAGATTGGCGATCCC |
| 16 | **pIJLK – seq – rv** | CGTGCTATGATCGACTGATG |
| 17 | **pIJ10257 – seq – fw** | AGATGGTTACCTCGCCTCTG |
| 18 | **pIJ10257 – seq – rv** | TCAGCGAGCTGAAGAAAGAC |
| 19 | **pEKEx2 – seq – fw** | GGAAAGCCACGTTGTGTCTC |
| 20 | **pEKEx2 – seq – rv** | GCCTCGTGAAGAAGGTGTTG |
| 21 | **pAN6 – seq – fw** | GATATGACCATGATTACGCCAAGC |
| 22 | **pAN6 – seq – rv** | CGGCGTTTCACTTCTGAGTTCGGC |

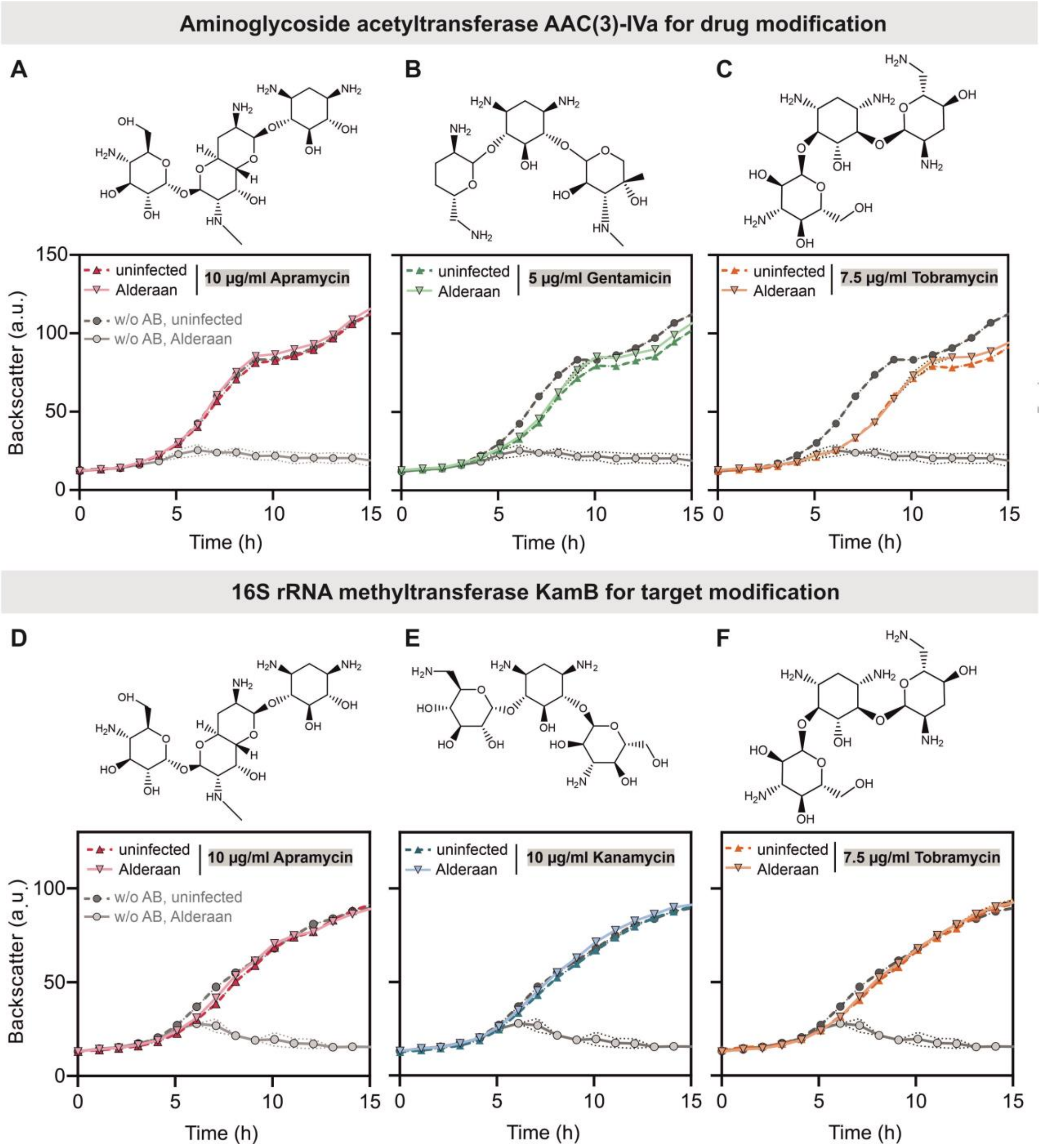


**Figure S1*: Inhibition of phage Alderaan infection by structurally-divergent aminoglycosides in aminoglycoside-resistant S. venezuelae* strains.** A-C) Growth assays of *S. venezuelae* ATCC 10712-*aac(3)-IVa* comparing infection with phage Alderaan in presence and absence of A) apramycin, B) gentamicin and C) tobramycin. Corresponding phage titers are presented in Figure 2G. D-F) Growth assays of *S. venezuelae* NRRL B-65442 - *kamB* comparing infection with phage Alderaan in presence and absence of D) apramycin, E) kanamycin and C) tobramycin. Corresponding phage titers are presented in Figure 2G.

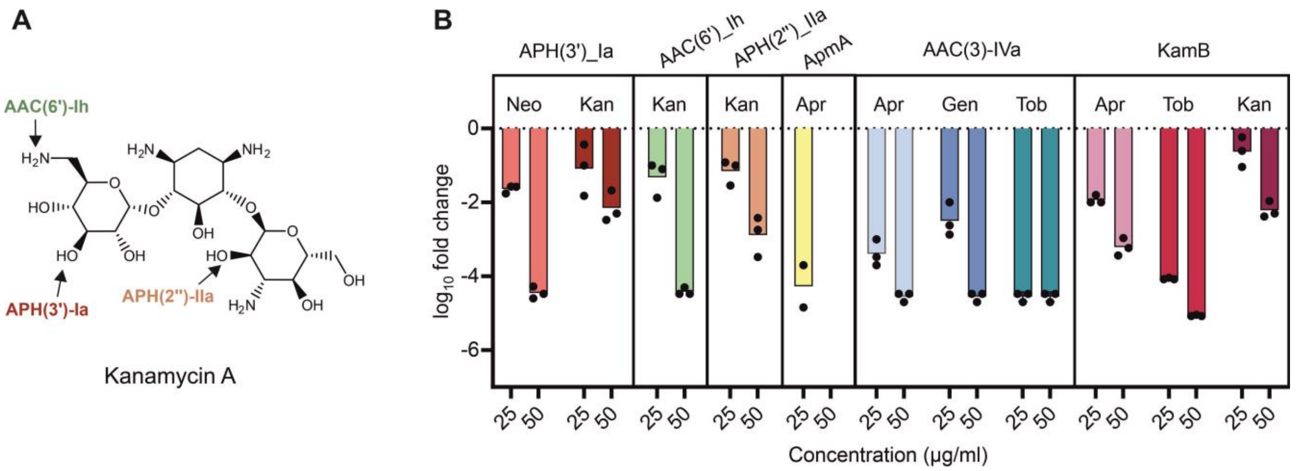


**Figure S2: Influence of different aminoglycoside modifications on infection of *E. coli* by phage λ.** A) Modification positions of selected kanamycin resistance genes. b) $log_{10}$ fold change in PFU/ml observed for different resistance gene - aminoglycoside combinations. Infection was conducted on LB double-agar overlay assays in biological triplicates. Plaques were counted after 16 h of incubation at 37°C. In case of ApmA-mediated resistance, 50 µg/ml apramycin did not allow proper growth of the bacterial lawn.

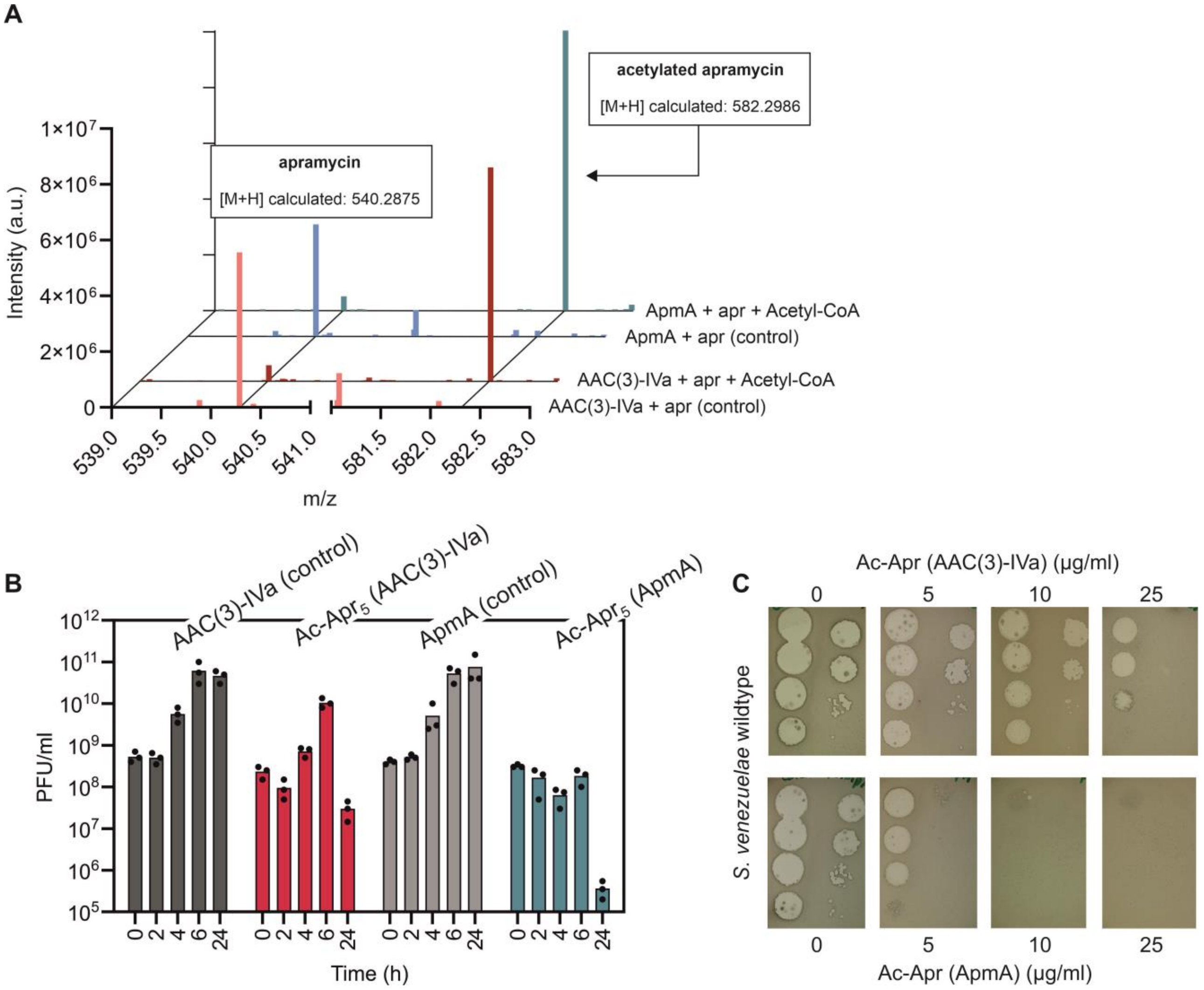


**Figure S3: Acetylation of apramycin with the two purified acetyltransferases AAC(3)-IVa and ApmA and its influence on phage infection dynamics.** (A) Mass spectrometry showing the successful in vitro acetylation of apramycin with the two purified acetyltransferases AAC(3)-IVa and ApmA. Control reactions without addition of the co-factor acetyl-CoA are used for comparison. Respective infection assays with acetylated apramycin are shown in Figure 2D and E. (B) Time-resolved quantification of Alderaan phage titers to infection assays shown in Figure 2D. (C) Double-agar overlays with *S. venezuelae* NRRL B-65442 wildtype and its phage Alderaan upon addition of acetylated apramycin.

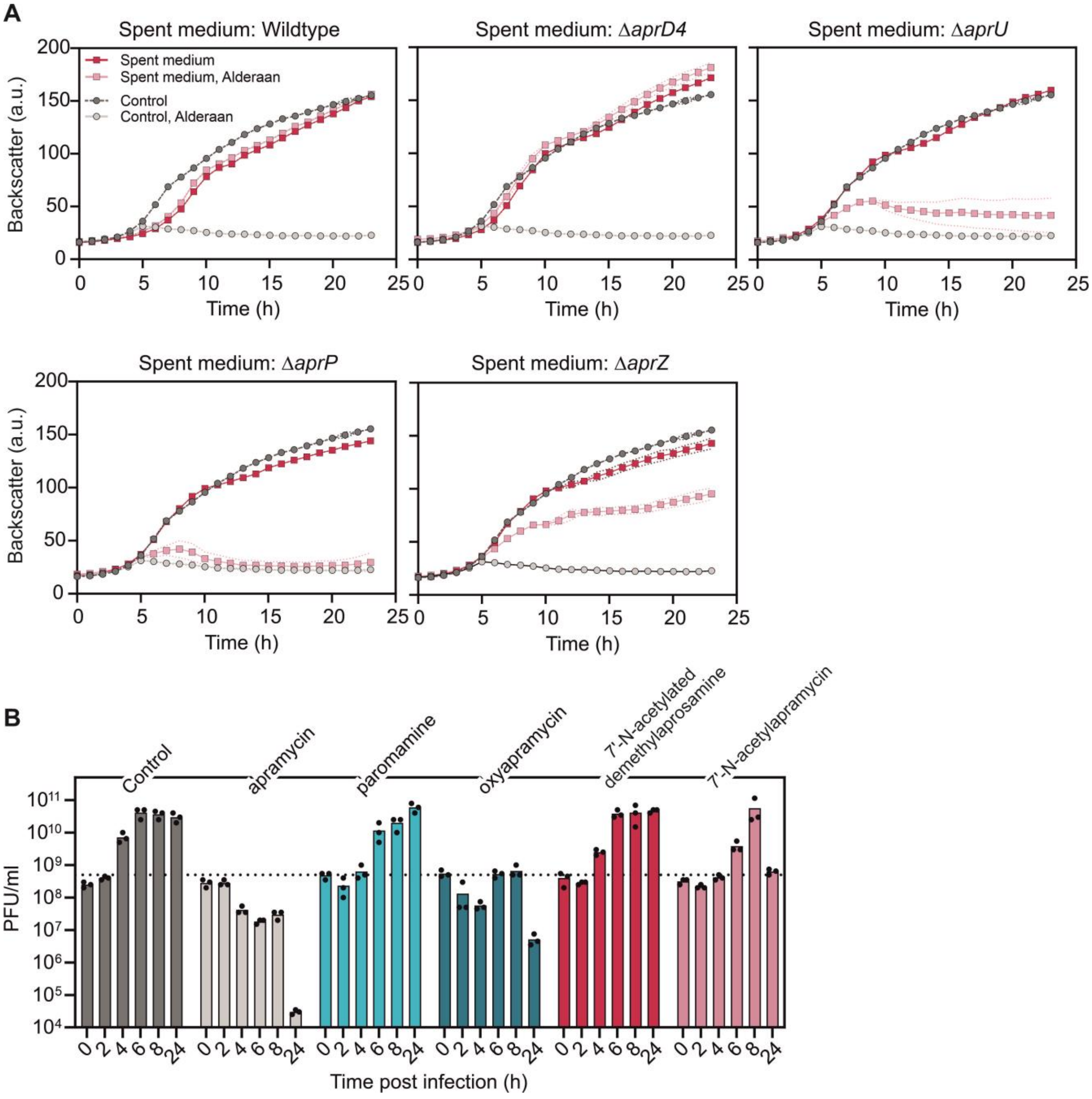


**Figure S4: Effects apramycin biosynthesis intermediates on Alderaan infection of *S. venezuelae* NRRL B-65442 - *kamB*.** A) Growth curves during Alderaan infection presence of spent medium from different *S. tenebrarius* strains. Corresponding infection assays are presented in Figure 4B. B) Phage amplification upon addition of different apramycin biosynthesis intermediates. Corresponding $log_{10}$ fold changes are shown in Figure 3C.
